# ProteinMPNN Recovers Complex Sequence Properties of Transmembrane β-barrels

**DOI:** 10.1101/2024.01.16.575764

**Authors:** Marissa Dolorfino, Rituparna Samanta, Anastassia Vorobieva

## Abstract

Recent deep-learning (DL) protein design methods have been successfully applied to a range of protein design problems, including the *de novo* design of novel folds, protein binders, and enzymes. However, DL methods have yet to meet the challenge of *de novo* membrane protein (MP) and the design of complex β-sheet folds. We performed a comprehensive benchmark of one DL protein sequence design method, ProteinMPNN, using transmembrane and water-soluble β-barrel folds as a model, and compared the performance of ProteinMPNN to the new membrane-specific Rosetta Franklin2023 energy function. We tested the effect of input backbone refinement on ProteinMPNN performance and found that given refined and well-defined inputs, ProteinMPNN more accurately captures global sequence properties despite complex folding biophysics. It generates more diverse TMB sequences than Franklin2023 in pore-facing positions. In addition, ProteinMPNN generated TMB sequences that passed state-of-the-art in silico filters for experimental validation, suggesting that the model could be used in *de novo* design tasks of diverse nanopores for single-molecule sensing and sequencing. Lastly, our results indicate that the low success rate of ProteinMPNN for the design of β-sheet proteins stems from backbone input accuracy rather than software limitations.

## Introduction

*De novo* protein design has promise in solving a broad range of problems in the fields of drug and vaccine development [1–2], biotechnology [3–4], and climate change [5–6]. Until recently, state-of-the-art protein design methods have relied on physics-based, statistical, or hybrid models such as Rosetta [7]. However, protein design with Rosetta is computationally expensive and requires structural biology expertise to generate protein models with new properties but native-like features that enable successful folding [8]. Recent deep-learning (DL) design methods enable faster, more accurate prediction of 3D protein folds [9–11] and the design of novel protein backbones [12–13] and sequences [14–15]. ProteinMPNN is a graph-based DL sequence design model that generates a protein sequence matching an input protein backbone. It consists of 3 encoder layers that encode the backbone coordinates of the input protein, followed by 3 decoder layers that predict a sequence in seconds and in an autoregressive manner [14]. It is a powerful tool that has been successfully applied to many protein design problems, including the *de novo* design of new folds [16], protein binders [17] and enzymes [18], and the redesign of native proteins [19]. However, DL design tools have met limited success for protein folds mainly composed of antiparallel β-sheets [20]. β-sheet secondary structures are challenging to design due to the presence of long-range interactions and the intrinsic tendency of β-strands to aggregate if the folding into the native state is not precisely controlled [21–23]. Moreover, both traditional and DL approaches have yet to fully meet the challenge of designing *de novo* membrane proteins (MPs) [24–27]. Despite MPs representing 20-30 % of the proteome [28–29] and the potential of MPs to provide a new diversity of scaffolds, chemistries, and functions for protein engineering, MP design is limited by the amount of available experimental data [24][30]. Traditional physics-based design methods rely on an energy function to estimate the total free energy of the sampled protein sequence and conformation [30–32]. These energy functions are trained on experimental high-resolution structures and thermodynamic stability data [30][33]. DL based protein sequence design methods learn to map structure to sequence, forgoing the need for energy function optimization, but require large datasets for training [14–15]. Yet, MPs represent only 2.6 % of structures in the Protein Data Bank (PDB) [34].

Of particular interest in MP design are transmembrane β-barrels (TMBs), which form pores through the lipid membrane and are widely used by the nanopore sequencing industry [3][35–37]. The *de novo* design of TMBs has the potential to transform single-molecule sequencing and sensing technologies by providing a wealth of new nanopores tailored to recognize and transport specific molecules and polymers [21][30]. However, TMBs represent a class of MPs that are especially difficult to generate *de novo* because they combine the challenges associated with β-sheet and MP design. Previous success in *de novo* TMB design utilized the reference Rosetta energy function manually refitted to design sequences with TMB-like properties [7][21]. The successful design of TMBs using this method elucidated properties of designed TMB sequences that are necessary for proper expression and folding *in vitro*, including the importance of negative design (local destabilization of sequences) to generate sequences with weak β-sheet propensity that slow β-sheet assembly [21][38]. These findings were validated experimentally by Martin et al. [22]. Though remarkable, this Rosetta method for TMB design requires significant expertise which could be alleviated by the use of DL in the process. DL tools have yet to be applied to the general design of MPs, but ProteinMPNN has been successfully used to design soluble analogues of integral MPs, suggesting that it may also be suitable for designing MPs [23].

Here, we conducted a comprehensive benchmarking analysis of ProteinMPNN [14], one of the latest structure-to-sequence DL models, in the context of TMB design. We characterized the ability of ProteinMPNN to capture global sequence properties of TMBs, to design diverse sequences, and to design high quality sequences that are likely to fold *in vitro*. We compared the performance of ProteinMPNN to that of the recently developed Rosetta MP-specific energy function, Franklin2023 [39].

## Datasets and Experimental Design

In contrast to the combinatorial ‘flexible-backbone’ design with Rosetta that iterates between sequence sampling and energy minimization of the entire structure [40] and which was used for the original TMB design, ProteinMPNN considers only the fixed-backbone input and incorporates a minimal amount of backbone noise (< 0.5 Å) [14]. Therefore, non-native features present in the input backbone that would otherwise be attenuated by flexible-backbone design will likely have a strong effect on ProteinMPNN performance. The definition of antiparallel β-sheet backbones for *de novo* design is particularly challenging by comparison to other secondary structures, due to the larger conformational space accessible to β-strands and to the presence of long-range interactions [21–23]. This difficulty is exemplified by the low expression rate of β-barrel folds in previous design attempts with ProteinMPNN [23]. We hypothesized that the low success rate of β-sheet fold designs generated with ProteinMPNN stems from the presence of non-native features in the input backbones rather than software limitations. To test our hypothesis, we assessed the performance of ProteinMPNN on *de novo* designed β-barrel backbones with different levels of refinement. We curated sets of backbones that have been successfully used as starting points for the *de novo* design of 8-strands transmembrane [21–22] and water-soluble β-barrels [22] with very similar structural properties (**Figure 1A**). These designs were generated through several iterations of Rosetta flexible-backbone design and backbone models (∼50 backbones/set) from three stages of the design pipelines were used as input backbones for ProteinMPNN: (i) the initial coarse-grained poly-valine (CG) backbones [41]; (ii) the medium-refined (MR) backbones obtained using one round of Rosetta flexible-backbone design; and (iii) the fully-refined (FR) designs selected for experimental characterization in the original studies [21–22] (**Figure 1B**).

**Figure 1.**
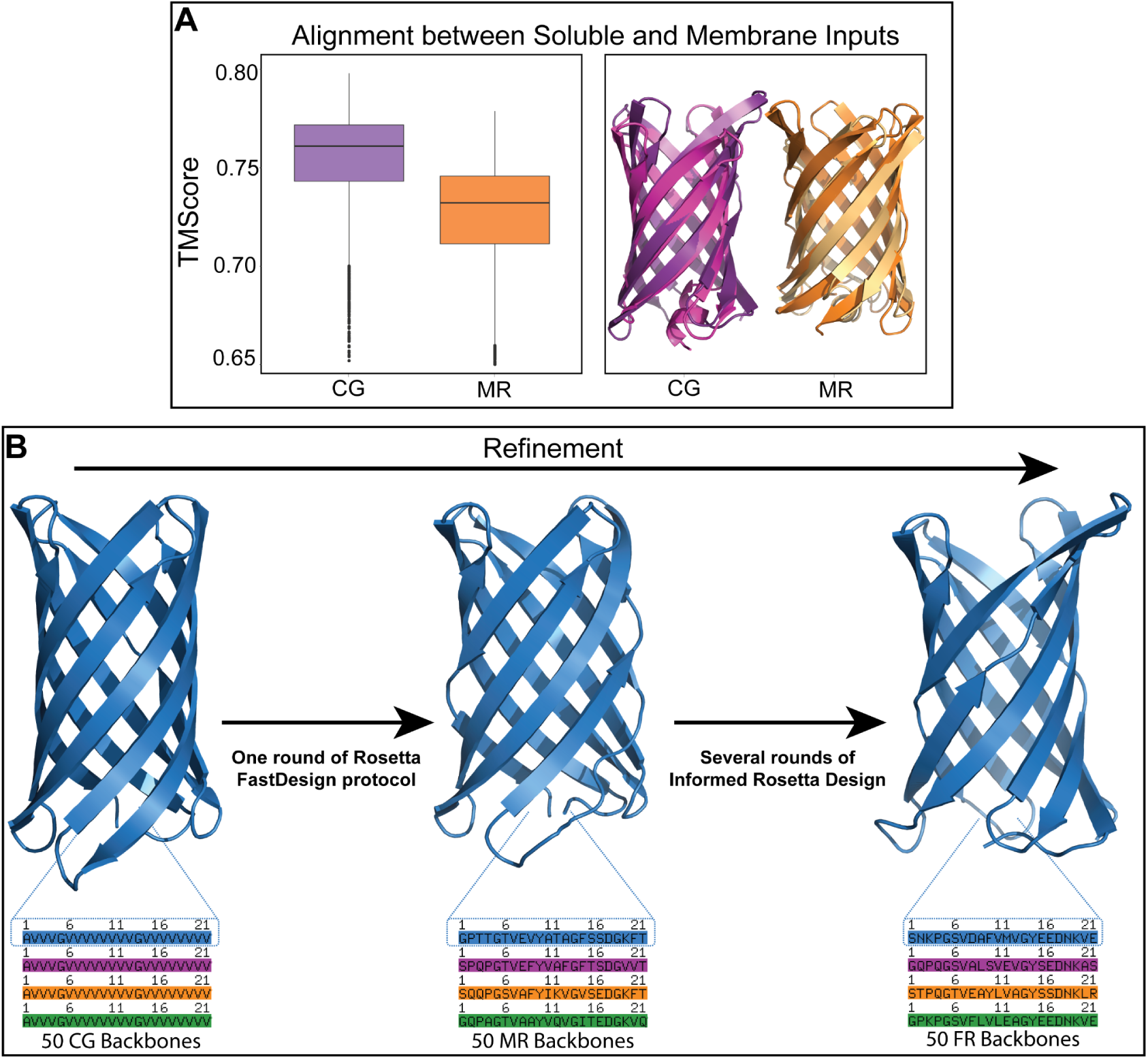
Soluble and transmembrane β-barrel models used for design with ProteinMPNN. (A) Soluble and transmembrane β-barrels share similar structural properties based on the TMscores [42] computed for all-to-all structure superposition (TMScore between soluble and transmembrane backbones (left) and aligned soluble and transmembrane backbones (right) from the CG and MR input sets). (B) A Rosetta pipeline was used to generate the three sets of input structures. The MR input set was generated from the CG input set using one round of the Rosetta FastDesign protocol (flexible-backbone energy minimization of sequence, backbone, and side chain conformations). The FR input set was generated from the MR input set using iterative rounds of the FastDesign protocol, selecting the most optimal designs for further refinement after each iteration.

### Sequence Property Recovery is Sensitive to Angstrom-level Changes of the Input Structure

We first sought to assess the ability of ProteinMPNN to recover global sequence properties for a given protein fold, using β-barrel backbones as test cases. While 8-strands transmembrane and water-soluble β-barrels have very similar structural properties (average TMscore [42] between CG and MR backbones of 0.75 and 0.73, respectively) (**Figure 1A**), the two folds drastically differ at the sequence level. Both folds feature the alternate polar/hydrophobic residue pattern characteristic of β-strands [21]. However, water-soluble β-barrels have large hydrophobic cores and hydrophilic surfaces while the hydrophobicity pattern is inverted in TMBs: the lipid-exposed surface is largely hydrophobic and the pore is water-accessible and hydrophilic [21]. To investigate the properties of sequences generated on similar β-barrel structures, we generated 10 sequences per input backbone using ProteinMPNN. The output sequences were evaluated based on their hydrophobicity patterns (calculated with the GRAVY hydropathy scale [43]; negative hydropathy values indicate polar properties, while positive hydropathy indicates hydrophobic properties), and TMB sequence patterns were compared to hydrophobicity patterns of native TMB sequences.

CG backbones of transmembrane and water-soluble β-barrels yielded very similar sequences with positive GRAVY scores for surface residues (2.18 and 2.12 for water-soluble and transmembrane β-barrels, respectively) and close to zero GRAVY scores for core residues (0.08 and -0.09 for water-soluble and transmembrane β-barrels) (**Figure 2A**). This result is consistent with the structural similarity between the input backbone sets (**Figure 1A**), which were generated using similar Rosetta pipelines [7][44]. The similar hydrophobicity patterns of the sequences might be reminiscent of the valine placeholders used at each residue position during backbone assembly. Surprisingly, MR inputs yielded ProteinMPNN sequences with clearly distinct properties and with hydrophobicity profiles matching water-soluble and transmembrane β-barrels (**Figure 2A**). Transmembrane β-barrel designs exhibited hydrophobic surfaces (average GRAVY score of 2.49) and hydrophilic cores (average GRAVY score of 0.02); water-soluble β-barrel designs exhibited largely hydrophilic surfaces (average GRAVY score of -1.92) and hydrophobic cores (average GRAVY score of 1.84). We quantified the structural change between the CG and MR β-barrel backbones (by comparing MR backbones to the corresponding parent backbone in the CG set) and found it to be quite small, with an average Root Mean Square Deviation (RMSD) of 1.12 and 1.45 Å for transmembrane and water-soluble models, respectively (**Figure 2B**). Hence, a single round of flexible-backbone design, restricted to native-like amino acid compositions characteristic of each fold, was sufficient to generate backbones yielding sequences with the expected hydrophobicity profiles. The sequence properties of designs generated from the FR backbones were similar to those from the MR set, indicating that further input refinement does not improve the ability of ProteinMPNN to accurately distribute hydrophobic and hydrophilic residues (**Supplementary** Figure 1). We compared the properties of the TMB sequences generated using ProteinMPNN to those generated with fixed-backbone design using the latest MP-specific Rosetta energy function Franklin2023 [39]. ProteinMPNN more accurately distributed hydrophobic and hydrophilic residues across TMBs (**Figure 2A**). A large majority of Franklin2023 TMB designs (82.9 %) had non-native sequence properties (hydrophobic pore-facing residues), potentially due to limitations of the geometric definition of the water-accessible pore environment [45]. Hence, if refined, accurate, and native-like backbones are provided as input, ProteinMPNN is able to recover global sequence properties of a given β-sheet fold with higher accuracy than a comparable fixed-backbone Rosetta protocol using an energy function trained specifically on MPs.

**Figure 2.**
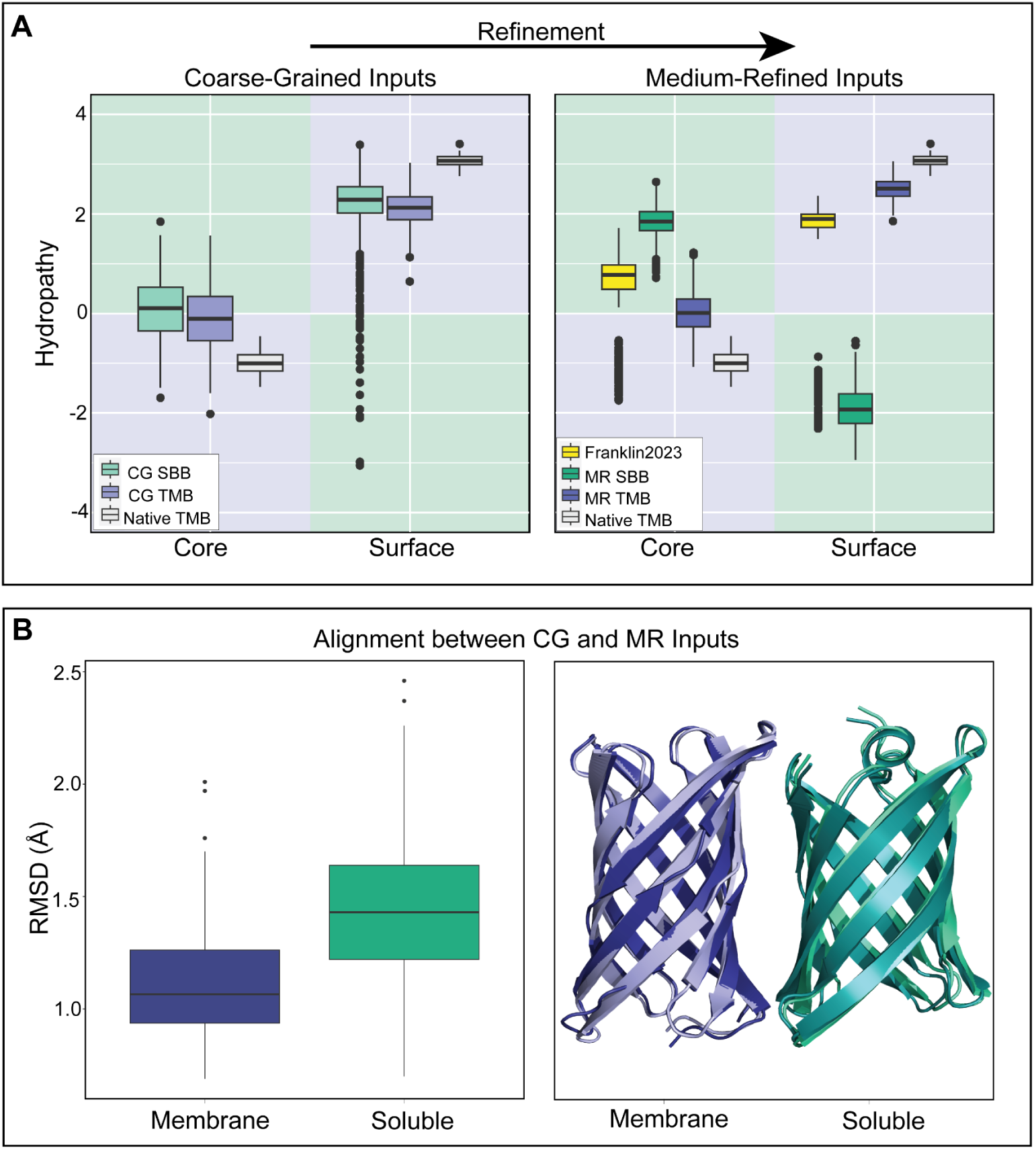
ProteinMPNN accurately recovers global sequence properties of water-soluble and transmembrane β-barrels with refined backbones. (A) CG inputs generate similar sequences for water-soluble and transmembrane backbones, while MR inputs correctly distinguish water-soluble and transmembrane inputs. TMB sequence patterns were also compared to native TMB sequence patterns. (B) One round of flexible-backbone design (between CG and MR backbone sets) results in structural changes of less than 1.5 Å for both transmembrane and water-soluble beta-barrels (RMSD between CG and MR backbones (left) and aligned CG and MR backbones (right) from the membrane and water-soluble input sets).

### Sequence Diversity of Designs

Since TMBs represent only 0.5 % of the training dataset of ProteinMPNN [14], we sought to evaluate the diversity of TMB sequences generated by ProteinMPNN to investigate potential overfitting of the model. We analyzed the diversity of the TMB sequences generated by ProteinMPNN and by Rosetta fixed-backbone design (with Franklin2023) using the FR transmembrane β-barrel backbones as input. The FR backbones yielded the best-scoring TMB models after several iterations of flexible-backbone sequence design [21–22]. They are therefore more “designable” [22] and more likely to incorporate native-like features enabling the design of successfully folding TMBs. We considered two “sequence diversity” criteria. First, we computed the sequence recovery (of surface-exposed, pore-facing, and all residues) as the percentage of identities between the generated sequences and the sequence for which each backbone was originally optimized with Rosetta (**Figure 3A**). The average sequence recovery of ProteinMPNN designs (72.05 %) was higher than the average sequence recovery of Franklin2023 designs (64.99 %). However, ProteinMPNN generates more diverse sequences in pore-facing positions (51.27 % sequence recovery) while the diversity of Franklin2023 designs is localized mainly on surface-exposed positions (43.84 % sequence recovery). Since the sensitivity and selectivity of a nanopore for a given molecule depends on the properties of residues lining the pore [3][20–21][35–37], ProteinMPNN is more likely than Franklin2023 to generate nanopores with diverse pore properties and functions. We next sought to investigate potential biases of the models by computing the recovery per amino acid across all core and surface residues (**Figure 3B**). In agreement with our previous observation, we found that ProteinMPNN was more likely to design diverse mutations changing the chemical properties of a residue, both on the surface and in the pore. In pore-facing positions, both polar-to-hydrophobic and hydrophobic-to-polar mutations were observed. Glycine and proline residues were strongly conserved, probably because of the constrained torsional space occupied by these amino acids in the Ramachandran plot [46]. In contrast, Franklin2023 designs frequently incorporate hydrophobic amino acids in place of polar residues in the pore. For example, Franklin2023 designs asparagine to arginine and glutamine to leucine with 80 % and 23 % frequency. While ProteinMPNN incorporated diverse mutations in surface-exposed positions, Franklin2023 was strongly biased against valine and alanine, mutating these residues to isoleucine and phenylalanine, respectively. Hence, despite the small number of TMBs included in the training dataset of ProteinMPNN, the DL model generates diverse and less biased sequences than Rosetta design with Franklin2023, particularly in pore-facing positions.

**Figure 3.**
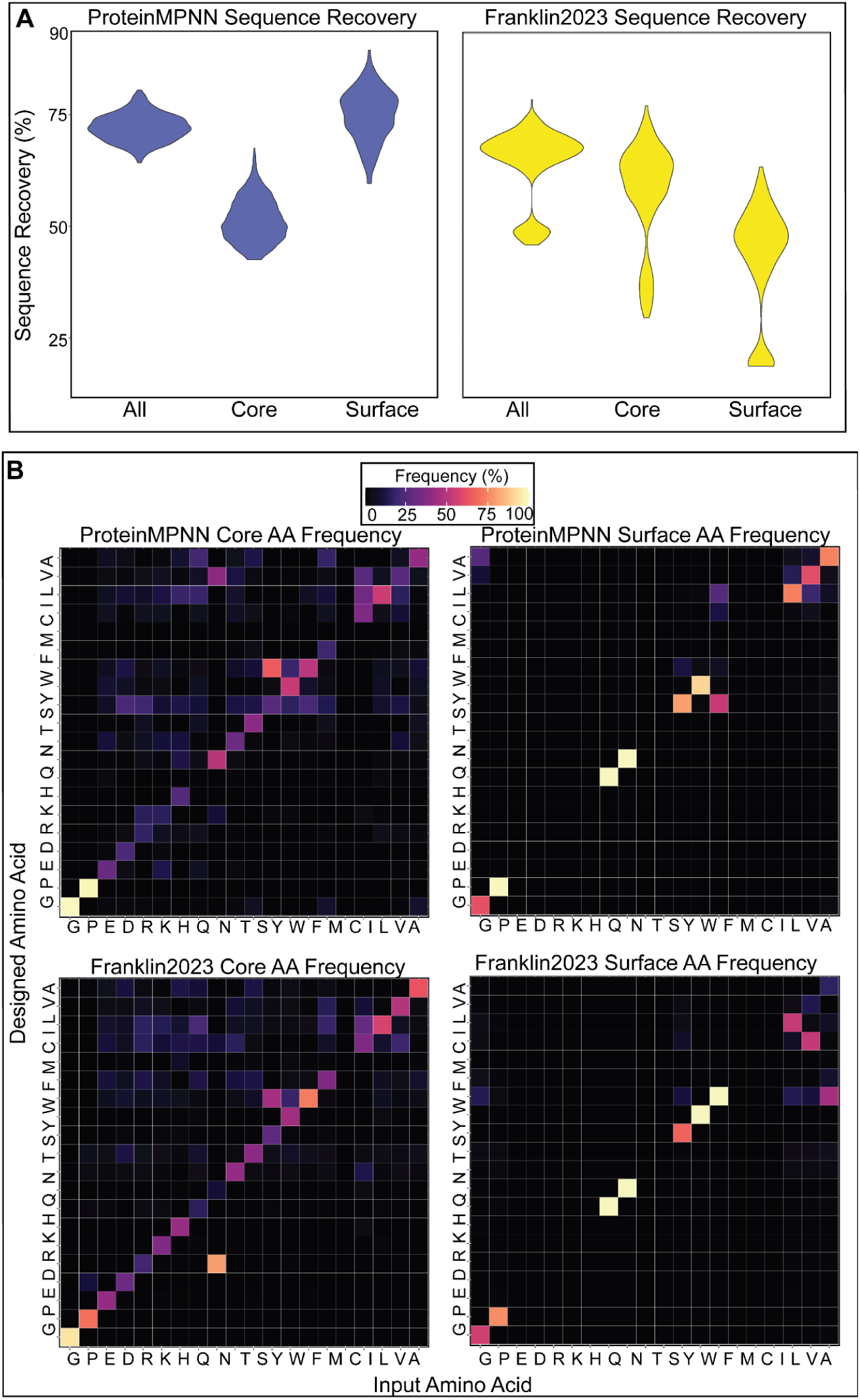
ProteinMPNN exhibits overall higher sequence recovery than Franklin2023, with a much higher surface residue sequence recovery but lower core recovery than Franklin2023 (A). Per amino acid recovery for surface and core residues shows potential biases of the models (B).

### Input Refinement Improves Design Quality

Because TMB nanopore design has great potential for nanopore biosensing technologies, we sought to determine whether the designed TMB sequences meet the quality criteria for experimental characterization (ordering criteria). Vorobieva and Martin et al. [21–22] previously established that TMB designs are more likely to fold *in vitro* if they have overall polar cores (sum of GRAVY scores across pore-facing residues < 0), have less than 50 % of residues predicted to fold into β-sheet secondary structure (< 50 % β-sheet propensity), and can be refolded *in silico* using AlphaFold2 from a single sequence (rather than from a multiple sequence alignment input). The calculations of core hydropathy and β-sheet propensity are detailed in the methods section. **Figure 4A** shows the distribution of β-sheet propensity and core hydropathy among the ProteinMPNN designs generated from the MR and FR input sets. We found that considerably more FR designs meet the empirical ordering criteria (16.6 % vs 5.1 %). This result is in agreement with our previously observed improvement in ProteinMPNN performance when refined and accurate TMB backbones are used as input. In comparison, 17.1 % of designs generated with Franklin2023 from the FR inputs have core hydropathy less than 0 and β-sheet propensity less than 50 % (**Figure 4B**). However, the designs generated with Franklin2023 that meet the ordering criteria had very low β-sheet propensity (< 30 %). We hypothesized that, although > 50 % β-sheet propensity yields TMB sequences that cannot fold, sequences with too low secondary structure propensity are equally unlikely to encode stable TMBs. To test this hypothesis, we used AlphaFold2 (ColabFold interface [47], single_sequence mode, 48 recycles, see methods) to predict 3D structures for the designed sequences. We evaluated the confidence of predictions (pLDDT) as well as the similarity (RMSD) between the predicted structures and the input model [9]. All of the MR (100 %) and almost all of the FR (94 %) designs pre-selected based on secondary structure propensity and core hydropathy were predicted with high confidence (pLDDT > 80) to fold into 3D structures very similar to the input model (RMSD < 2 Å) (**Figure 4**). None of the pre-selected Franklin2023 designs were predicted to fold into the desired β-barrel structure, validating our hypothesis (**Figure 4B**). Lastly, we assessed the ability of the methods to recover TMB sequences from native PDB structures, a common benchmark of protein design (**Supplementary** Figure 2). We found that ProteinMPNN had higher native sequence recovery (35.91 %) and generated sequences better encoding TMB structures (1.79 Å RMSD) than Franklin2023 (4.67 Å RMSD and 26.74 % sequence recovery).

**Figure 4.**
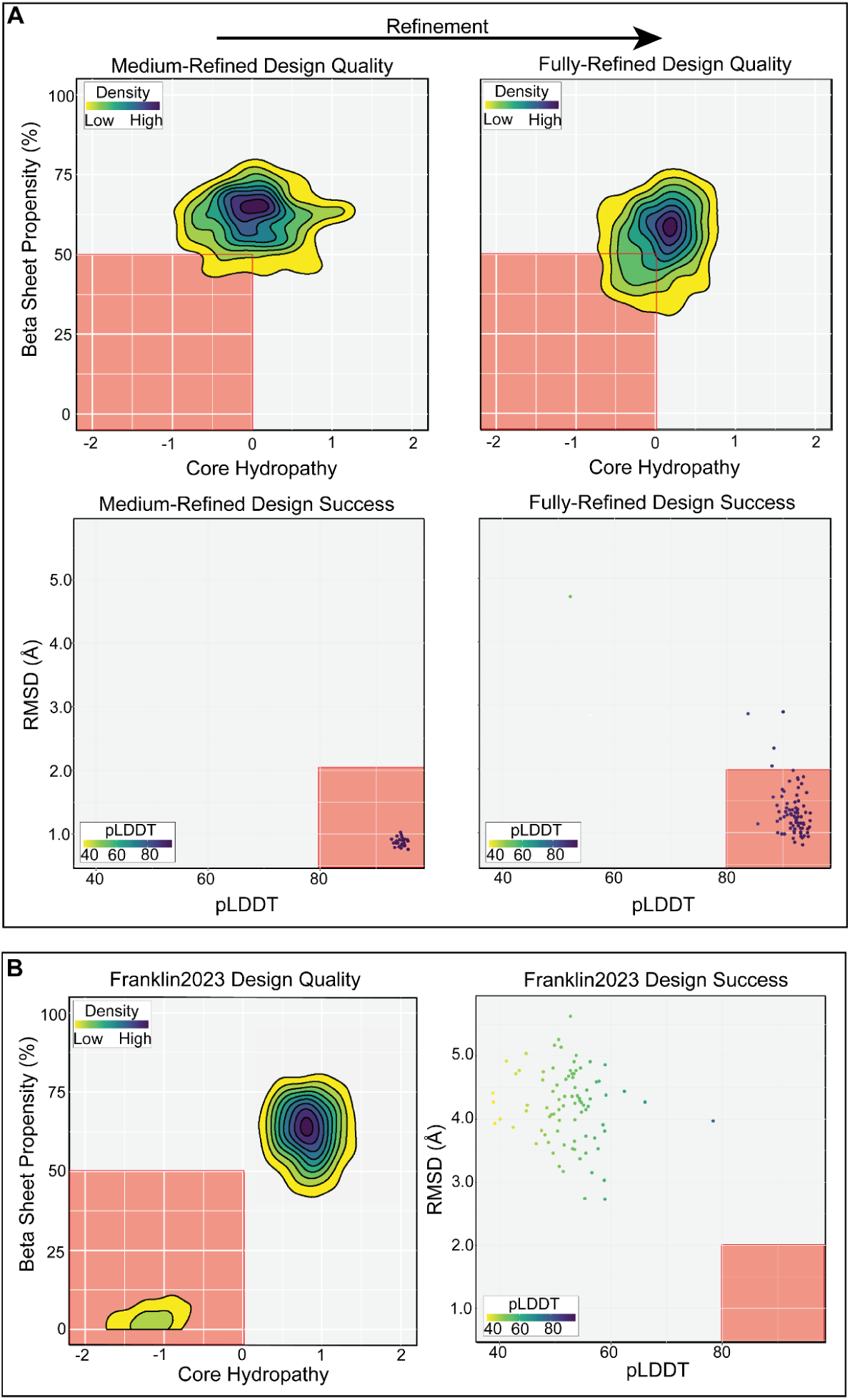
Measured ordering criteria and foldability criteria for ProteinMPNN (A) and Franklin2023 (B) designs. Design quality plots show the β-sheet and core hydropathy distribution of designs and design success plots show the RMSD of folded designs to the model vs the confidence of AlphaFold2 predictions (pLDDT).

## Discussion

ProteinMPNN is a powerful protein sequence design tool that has been applied to many different protein design tasks and applications. It is therefore critical to understand how its performance varies as a function of the quality of the input models and how it performs in design tasks of challenging protein folds. Here, we put ProteinMPNN to the challenge of designing TMBs, a difficult MP fold formed mainly from β-sheets. By performing a comparative study on very similar backbones used to design 8-strands water-soluble and transmembrane β-barrels, we show that ProteinMPNN is highly sensitive to angstrom-level structural differences in the input models. It is able to design sequences with the desired global properties (secondary structure and hydropathy) when presented with input backbones refined with atomistic details (as opposed to unrefined coarse-grained backbones), accurately placing hydrophobic and hydrophilic residues to generate realistic water-soluble and transmembrane β-barrels sequences. This suggests a high level of sensitivity to “backbone memory”. Yet, sequences generated with ProteinMPNN are quite diverse, with many sequences sharing less than 70 % sequence identity to inputs (a common cut-off for the selection of *de novo* designs). Moreover, amino acid substitutions generated by ProteinMPNN were less biased than those sampled with Rosetta fixed-backbone design and the membrane-specific energy function Franklin2023.

A major challenge of TMB design is the complexity of the folding biophysics. On the one hand, successful TMB design requires negative design to introduce local secondary structure frustration. On the other hand, the sequence should still have sufficient β-sheet propensity to encode a stable β-barrel structure. To design TMBs *de novo*, this fine balance is controlled by selecting sequences that feature limited predicted β-sheet propensity and yet are predicted to fold into a β-barrel structure by AlphaFold2. Only ProteinMPNN was able to generate sequences that simultaneously meet both selection criteria. Designs generated using Franklin2023 either had high predicted β-sheet propensity or could not be folded by AlphaFold2.

From this benchmark, we concluded that with accurate input refinement, ProteinMPNN is capable of successfully designing both water-soluble and transmembrane β-barrel protein folds with high diversity of pore-facing residues that are likely to fold *in vitro*. This indicates that ProteinMPNN could be used in the *de novo* design of nanopores for single-molecule sequencing and sensing. Our benchmark also produces evidence to suggest that ProteinMPNN is capable of designing accurate and foldable sequences for even the most difficult protein folds given refined, accurate, and native-like input structures. One limitation, however, is that ProteinMPNN is constrained to fixed-backbone design, while more traditional methods like Franklin2023 are capable of flexible-backbone design.

## Methods

### Input Set Refinement

The input backbone sets were generated using a Rosetta design pipeline (**Figure 1B**). For both soluble and transmembrane β-barrels, a set of 50 centroid poly-valine backbones were first curated, with glycine residues placed to obtain the characteristic β-barrel structure. This set comprised the coarse-grained (CG) input set. The medium-refined (MR) input set was generated from the CG input set by performing one round of the Rosetta flexible-backbone FastDesign protocol, which searches for low energy sequence and structural conformations of a given input. The fully-refined inputs were then generated from the MR input set by several rounds of the Rosetta flexible-backbone FastDesign protocol. At each round and for each MR structure, several optimized protein structures were generated. At the end of each design round, the most energetically favorable designs were chosen as inputs for the next round, akin to evolutionary selection. In total, each input set consists of roughly 50 protein backbones.

### ProteinMPNN Design

With ProteinMPNN, we generated 10 sequences for every input backbone, with a sampling temperature of 0.1. We allowed free design of the sequences, with the exception of the residues in the loops of the β-barrels, which are highly conserved in native β-barrel structures.

### Franklin2023 Design

We generated 10 sequences for every input backbone using the Franklin2023 energy function with the Rosetta FastDesign protocol. We kept the loop regions fixed, and constrained the optimization to a fixed-backbone optimization.

### Hydropathy Calculation

To calculate the hydropathy of the sequences, we used the GRAVY hydropathy scale. For all residues in a region of interest (e.g. core or surface), we summed the GRAVY hydropathy score across the residues in that region. Negative GRAVY scores indicate a polar region, while positive GRAVY scores indicate a hydrophobic region.

### Sequence Diversity

The sequence recovery of designs was calculated as the percentage of identities between the generated sequences and the sequence for which each backbone was originally optimized with Rosetta. Sequence recovery per amino acid was determined by calculating the frequency of each input amino acid to design into any of the 20 amino acids in the designed sequences. For example, the percentage of alanine to phenylalanine residues is calculated by calculating the number of instances an alanine residue in an input sequence was designed into a phenylalanine in a designed sequence.

### AlphaFold2

The AlphaFold2 ColabFold version was used to predict protein structures [47]. ProteinMPNN and Franklin2023 design sequences were folded using AlphaFold2 with no MSA and no template. The number of recycles through the model was 48, consistent with the protocol proposed by Martin et al. [22], and 5 structures were generated per sequence. The structure AlphaFold2 predicted as the top ranking structure (by pLDDT) was the structure used for analyses.

### β-Sheet Propensity

To calculate the β-sheet propensity of the TMBs, we used the RaptorX secondary structure prediction program [48]. RaptorX predicts secondary structure from amino acid sequences, predicting the secondary structure of each residue in a sequence. We used a reference sequence from a native TMB with assigned secondary structures per residue. We compared the RaptorX predicted secondary structure from each design sequence to the reference sequence. The β-sheet propensity was then calculated by taking *n*_β_/*n_ref_*\*100, where *n_β_* is the number of correctly predicted β-sheet residues in the designed sequences and *n_ref_* is the number of β-sheet residues in the reference sequence.

### Structure Alignment

To align protein structures, we calculated Root Mean Square Deviation (RMSD) if the structures contained the same number of residues (in the case of aligning structures from the same environment) and TMscore if the structures contained differing number of residues (in the case of aligning soluble structures to membrane structures).

### Native Designs

We curated a set of 8 native TMB structures to calculate native sequence and structure recovery (PDB codes: 1QJP, 2LHF, 2VDF, 2X27, 2X55, 3DZM, 3QRA, 4RLC).

## Author contributions

MD and AAV conceived of this project as part of the RosettaCommons REU program. MD wrote the ProteinMPNN design and AlphaFold2 pipeline and generated and analyzed all ProteinMPNN designs. RS generated the Franklin2023 designs and MD analyzed Franklin2023 designs. AAV generated the input backbones using a Rosetta pipeline. MD wrote the manuscript with input from AAV and RS.

## Acknowledgment

This work was supported by the RosettaCommons REU program (to MD), core funding from the Flanders Institute of Biotechnology (VIB) (to AAV), the Fonds voor Wetenschappelijk Onderzoek - Vlaanderen (FWO) (grant FWOAL1092). The resources and services used in this work were provided by the VSC (Flemish Supercomputer Center), funded by the Research Foundation - Flanders (FWO) and the Flemish Government (MD, AAV). The resources used to generate Franklin2023 designs were supported by the National Institute of Health through grant R35GM141881-03 (to RS) and the facilities at the Advanced Research Computing at Hopkins (ARCH), which is supported by the National Science Foundation (NSF) grant number OAC1920103 (to RS). We thank Prof. Jeff Gray for insightful discussions and comments during formation of the project.

## Data availability

The python and R scripts used for data analysis are available from the TMB Design GitHub repository.

---

**Supplementary Figure 1.**
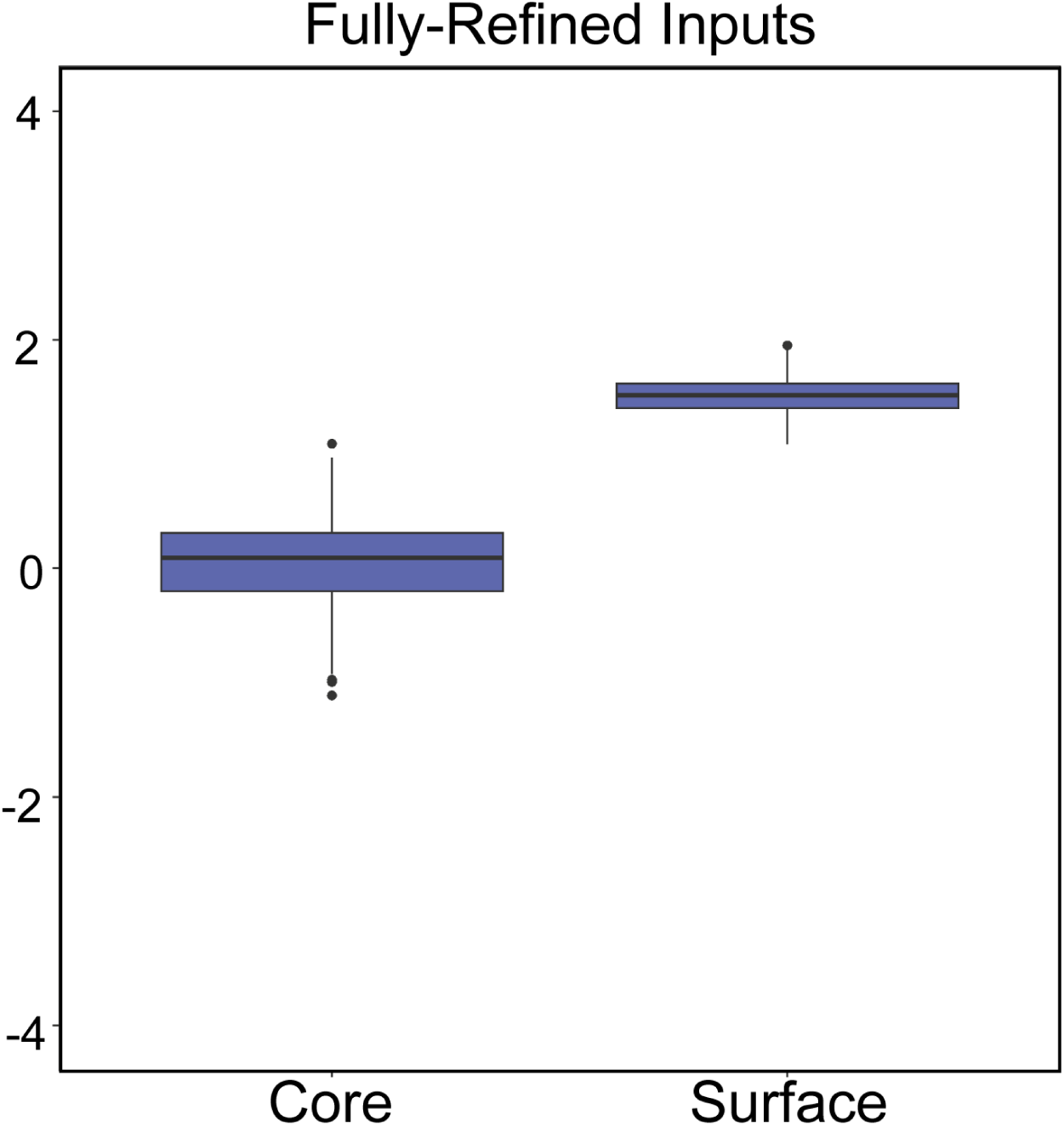
The distribution of hydrophobic and hydrophilic residues in designs generated using ProteinMPNN on the fully-refined input set is similar to the distribution observed with designs generated using ProteinMPNN on the medium-refined input set.

**Supplementary Figure 2.**
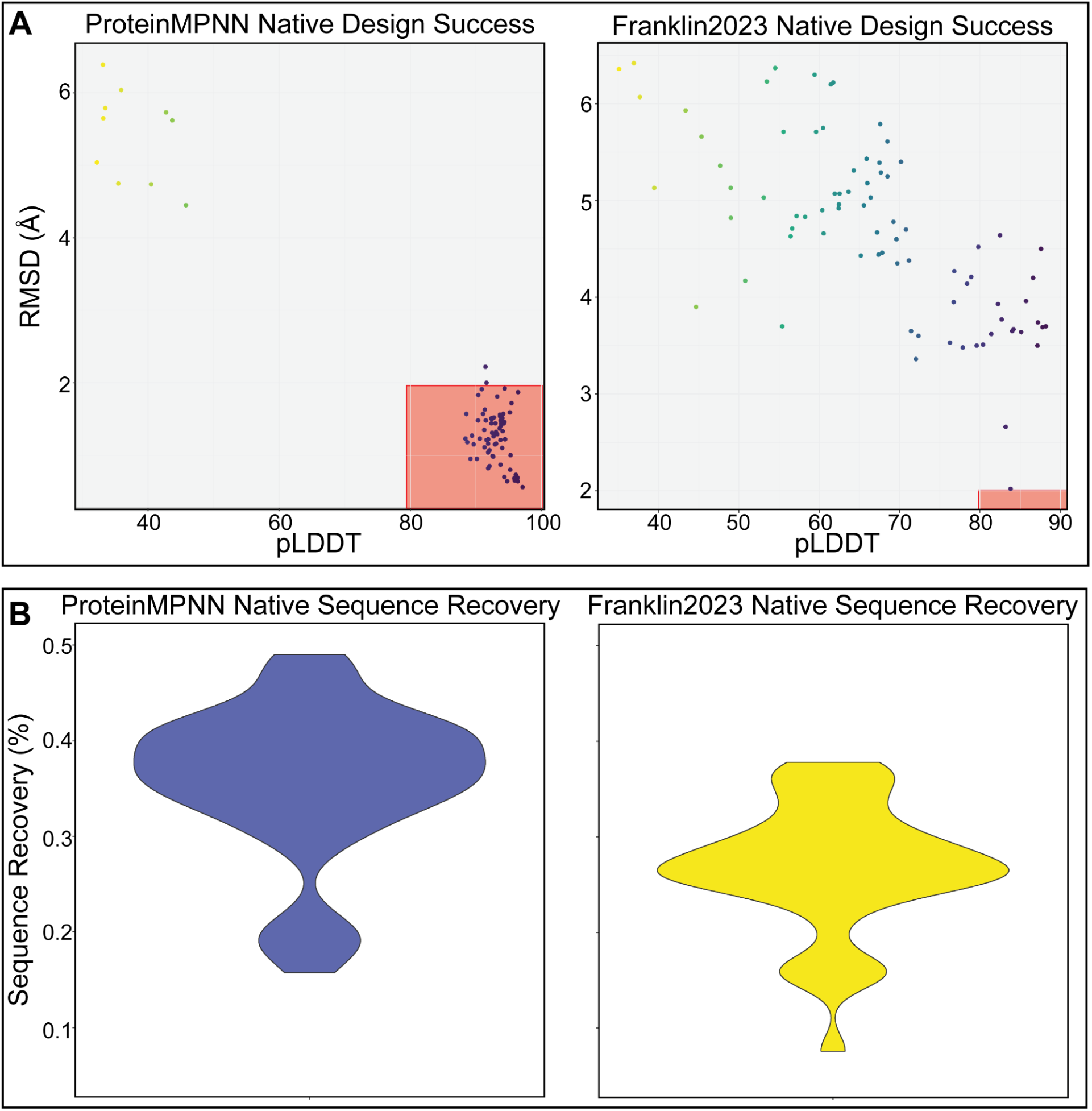
Recovery of native structures and sequences using ProteinMPNN and Franklin2023. ProteinMPNN designs have higher structure recovery (A) and sequence recovery (B) than Franklin2023 designs.

## Notes

https://github.com/marissadolorfino2024/ProteinMPNN-TMB-Design.git

### Competing Interest Statement

The authors have declared no competing interest.

### Summary of Updates

Added author after affiliations were sorted.

