## Supplementary figures and images for "ProteinMPNN Recovers Complex Sequence Properties of Transmembrane β-barrels"

### Supplementary Figure 1

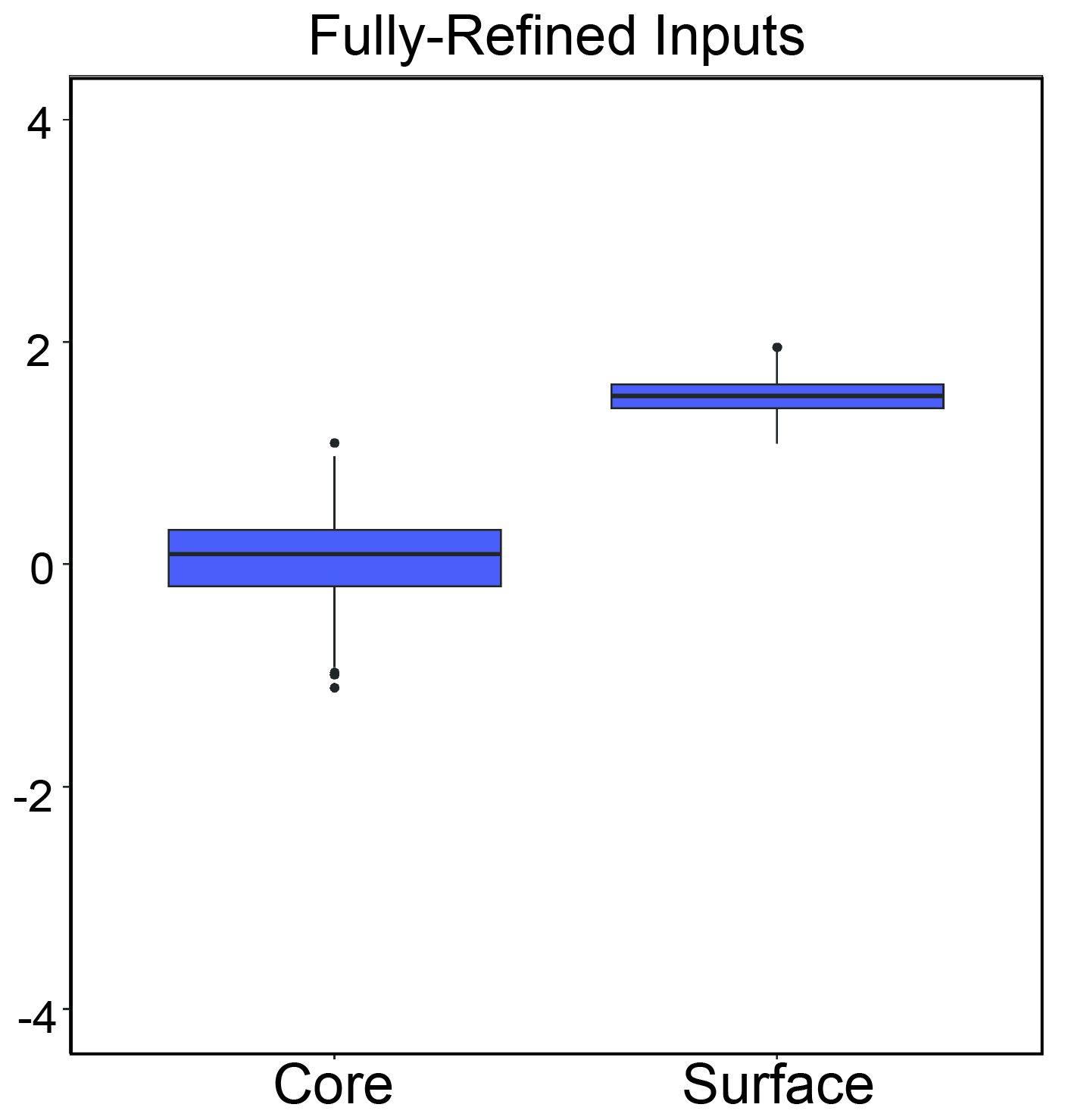

### Supplementary Figure 2

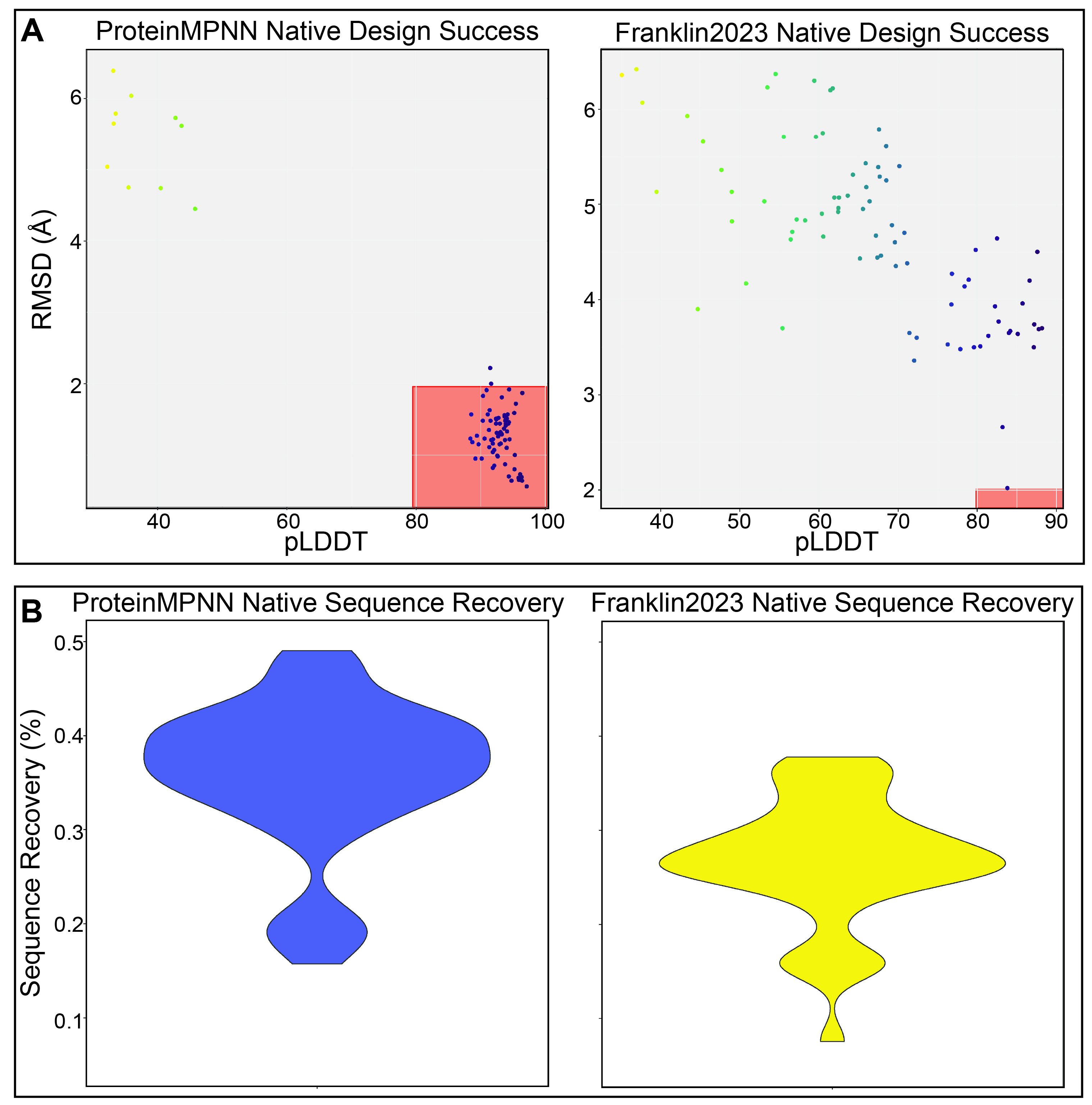
